# Structure modeling hints at a granular organization of the Vertebrate Golgi ribbon

**DOI:** 10.1101/2021.08.30.458089

**Authors:** Karen M. Page, Jessica J. McCormack, Mafalda Lopes-da-Silva, Francesca Patella, Kimberly Harrison-Lavoie, Jemima J. Burden, Ying-Yi Bernadette Quah, Dominic Scaglioni, Francesco Ferraro, Daniel F. Cutler

## Abstract

Vertebrate cells display a specific Golgi apparatus architecture, known as the “ribbon”, where the functional subunits, the mini-stacks, are linked into a tridimensional network. The importance of the ribbon architecture is underscored by evidence of its disruption in a host of diseases, but just how it relates to the biological Golgi functions remains unclear. Are all the connections between mini-stacks functionally equal? Is the local structure of the ribbon of functional importance? These are difficult questions to address, due to the lack of a secretory cargo providing a quantifiable readout of the functional output of ribbon-embedded mini-stacks. Endothelial cells produce rod-shaped secretory granules, the Weibel-Palade bodies (WPB), whose von Willebrand Factor (VWF) cargo is central to hemostasis. In these cells, the Golgi apparatus exerts a dual control on WPB size at both mini-stack and ribbon levels. Mini-stack dimensions delimit the size of VWF ‘boluses” while the ribbon architecture allows their linear co-packaging at the *trans*-Golgi network generating WPBs of different lengths. This Golgi/WPB size relationship lends itself to mathematical analysis. Here, different ribbon structures were modeled and their predicted effects on WPB size distribution compared to the ground truth of experimental data. Strikingly, the best-fitting model describes a Golgi ribbon made by linked subunits corresponding to differentially functioning monomer and dimer ministacks. These results raise the intriguing possibility that the fine-grained structure of the Golgi ribbon is more complex than previously thought.

## Introduction

In multicellular organisms, such as plants and most animals, the Golgi apparatus is a multi-copy organelle with subunits, the mini-stacks, scattered throughout the cell, often in close contact with endoplasmic reticulum exit sites (1–3). In contrast, in vertebrates mini-stacks link to each other into a tridimensional network, the Golgi ribbon (4–6). This Golgi arrangement is highly dynamic, undergoing regulated disassembly/reassembly during mitosis, directed migration and polarized secretion (6). The ribbon also disassembles in response to membrane depolarization, raised intracellular calcium, viral infection, autophagy induction and genotoxic insults (7–11). The importance of the Golgi ribbon in the physiology of vertebrate cells is further underscored by accumulating observations showing that a host of human diseases, especially neurodegenerative, exhibit disruption of ribbon morphology, collectively referred to as “Golgi fragmentation” (12–14). Nevertheless, the biological functions of the Golgi ribbon remain unclear (15), likely for two main reasons. First, understanding of the contributions of individual components of the complex molecular machinery involved in ribbon formation and maintenance is incomplete (16–28). Second, a lack of cellular systems where the detailed structure of the ribbon is intimately linked to a quantifiable biological function has prevented much experimental probing of the effects of this Golgi organization on a measurable biological output. Recently, this latter situation changed. Weibel-Palade bodies (WPBs) are endothelial-specific uniquely rodlike secretory granules with lengths varying over a 10-fold range (0.5-5μm). WPB length reflects the dual-level structural arrangement of the vertebrate Golgi apparatus. The major WPB cargo, the pro-hemostatic secretory protein Von Willebrand Factor (VWF), is molded into boluses during its *cis-trans* passage through the functional subunits of the Golgi apparatus, the ministacks. Given the narrow size range of the mini-stacks, these VWF boluses also occur within a similar small range (29). At the Golgi exit site, the continuous structure of the *trans*-Golgi network allows co-packaging of adjacent VWF boluses into a linear array that drives the formation of nascent WPBs in an AP1 complex-dependent process (29–31). The ribbon architecture of the Golgi apparatus thus determines the size of WPBs, which reflects the number of boluses co-packaged at biogenesis (**Figure 1A**). For this reason these VWF boluses were dubbed “quanta” (29).

**Figure 1.**
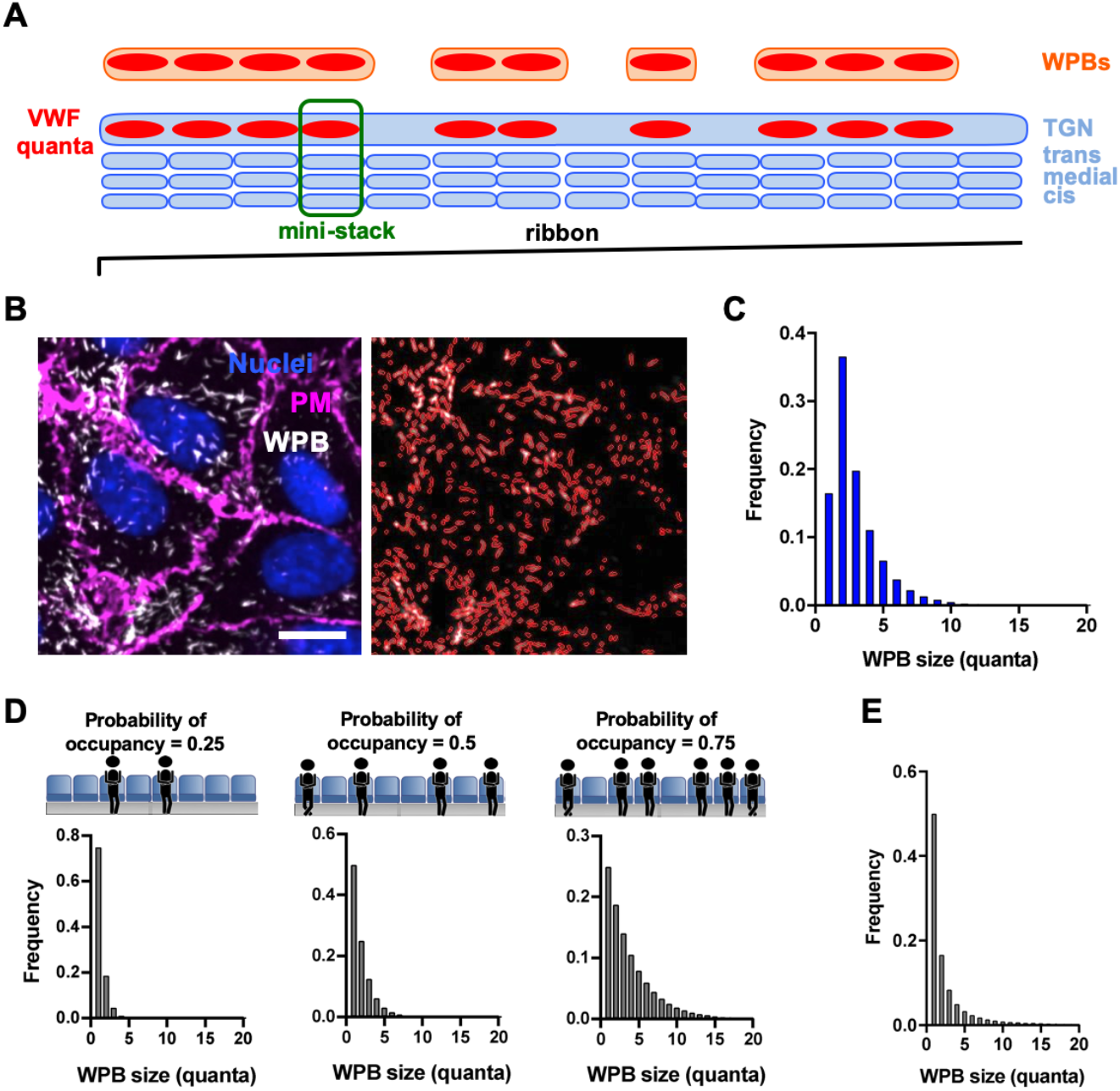
Modeling the Golgi ribbon as a linear array of mini-stacks. (A) Schematic representation of WPB biogenesis at the Golgi. In the context of the Golgi ribbon, the size of mini-stacks (highlighted in the green box) delimits that of VWF transiting quanta (red); at the Golgi exit site, the continuous lumen of the TGN allows co-packaging of adjacent quanta into nascent WPBs (orange). (B) On the left, a micrograph showing HUVECs immuno-stained for the plasma membrane (PM, with an anti-VE-cadherin antibody) and WPBs (with antibody which visualizes processed VWF at the Golgi and in WPBs) and counterstained for DNA (Nuclei) with Hoechst. On the right, WPBs have been segmented for quantification. Scale bar: 10 μm. (C) Measured WPB sizes (Figure S1A) were quantized by dividing the length of each organelle by the median quantum size (see text for details) and rounding to the nearest integer. (D) WPB size distributions as calculated from equation 1 at the indicated values of *p*, the probability of mini-stack occupancy by VWF quanta. (E) WPB size distribution calculated from equation 2, where *p* follows a probability distribution to account cell-to-cell variability in VWF expression.

WPB size determination can be summarized thus:

i. Mini-stack dimensions determine the size of VWF quanta. Changing the size of the mini-stacks also changes VWF quantum size and, as a consequence, WPB length distribution.
ii. The ribbon allows for VWF quanta occupying adjacent mini-stacks to be co-packaged into the same secretory granule. Unlinking of the ribbon results in WPB size distributions shifting to shorter lengths.
iii. Mini-stack occupancy by VWF quanta occurs by chance. The level of VWF expression controls the fraction of occupied mini-stacks, i.e., the number of VWF quanta formed. As a consequence, reducing VWF expression lowers the number of quanta within a Golgi ribbon, though not their size, resulting in fewer and shorter WPBs.

Statements *i-iii* are based on experimental evidence (29, 32).

This model of secretory granule biogenesis therefore invokes a key role for Golgi architecture, against which the level of VWF cargo controls both number and size of WPBs. Based on these premises, it is possible to mathematically model the effects of different organizations of the Golgi ribbon on WPB size and compare such simulations with experimental data obtained using a morphometric analytical workflow we developed to count numbers and measure the size of WPB (29). The best-fitting model obtained with this approach indicates that the WPB size distribution observed in endothelial cells could be generated by a Golgi organization in which the ribbon physically integrates mini-stacks functionally behaving as monomers and dimers. Therefore, mathematical modeling of a system where the relationship between structure and function of the Golgi ribbon is clear and quantifiable raises the intriguing possibility that vertebrate mini-stack subunits are functionally - and presumably structurally - different.

## Results

### Modeling the Golgi ribbon

The experimental data against which models were tested were generated with automated high-throughput morphometric analyses of cultured primary human umbilical vein endothelial cells (HUVECs). Immunostained WPBs were segmented to extract their length (**Figure 1B**). Our model of WPB biogenesis (statements *i-iii*, above) invokes a central role for the ministack size in constraining the size of VWF quanta. In fact, the median diameter of a Golgi mini-stack in HUVECs as measured in electron micrographs is 0.612 μm, which correlates with the median length of a VWF quantum (0.576 μm), determined by super-resolution microscopy (29). In order to simplify mathematical modeling, we equated mini-stack length (*I*) to the median size of a VWF quantum and expressed WPB lengths in multiples of *l* (by rounding up to the nearest multiple), thus transforming a distribution of WPB size from length (**Figure S1A**) to number of quanta (Q), or size class (i.e., 1Q-WPB, 2Q-WPB, etc.; **Figure 1C**).

To model how the Golgi generates this observed WPB size distribution, we started with the simplest assumptions:

a. the Golgi apparatus is a single, very long (effectively infinite) ribbon composed of linearly interlinked, identical mini-stacks, each of fixed length *l*.
b. mini-stacks are occupied at random with a probability *p* by a VWF quantum.
c. c) VWF quanta occupying adjacent mini-stacks will be packaged together into the same WPB. If, for example, five contiguous mini-stacks are occupied, with empty mini-stacks at both sides, a WPB of length five quanta (a 5Q-WPB) will form. *Note. The infinite ribbon assumption is made to avoid (i) cell-dependent variations; (ii) to reflect, like the WPB distribution, a cell population; and also (iii) to simplify probability calculations*.

Given these assumptions, by denoting with *N* the number of quanta in a WPB (i.e., its size) and with *p* the probability of VWF quanta occupancy of mini-stacks across the Golgi ribbon, we can calculate the expected distributions of WPB lengths when *p* varies (**Figure 1D**) with the following equation.

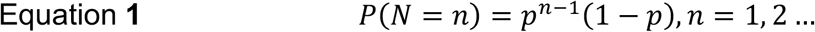

Independently of the value of *p*, all WPB length distributions predicted with this model display a mode at *N* = 1 (**Figure 1D**) and therefore do not simulate the experimental data that show a mode at *N* = 2 (**Figure 1C**). Furthermore, the 2Q-WPB frequency in the experimental distribution, 0.366, is much higher than expected by the modeling at any value of *p*; in fact, the greatest proportion of 2Q-WPBs predicted by the model is 0.25 for *p* = 0.5 (**Figure 1D,** middle panel).

### Variable mini-stack occupancy

We next considered whether the level of mini-stack occupancy might affect the relative frequency of the WPB size classes. Assuming that the rates at which WPBs are formed and that the rates at which they disappear from the cells, either because of exocytosis or degradation, are size-independent, one would expect cells with mini-stack occupancies close to 0.5 to have the most WPBs, since at very low occupancy there should be fewer WPBs, most of which should be 1Q-WPB, whereas at very high occupancy there should also be few WPBs but they should be close to the length of the Golgi ribbon.

Since cell-to-cell variability in the levels of VWF synthesis is observed in endothelial cells (33) and the HUVECs used in our lab are pooled from multiple donors, the probability of occupancy of Golgi mini-stacks by VWF quanta varies from cell to cell (**Figure 1B** and **Figure S1B**). Trying to explain the observed WPB size distribution, captured from a cell population, by assuming a single value of *p* is therefore an oversimplification. We thus modified the predictive model by posing that *p* follows a probability distribution. If *p* is uniformly distributed between 0 and 1 within a population of endothelial cells, we obtain equation **2**:

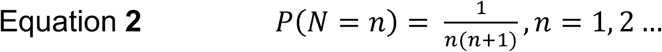

For every value of *p* this model predicts that the proportion of 2Q-WPBs is always ≤ 0.25 and the proportion of WPBs of length *N* strictly decreases with *N*. Therefore, even accounting for VWF expression variability does not explain the high proportion of measured 2Q-WPBs (**Figure 1E**).

### Fitting 2Q-WPB frequency

Our initial model is based on assumptions **a-c**. Assumptions **b** and **c** derive from our model of WPB size determination (statements *iii* and *ii*, respectively), which are experimentally supported. Assumption **a,** a very long, single ribbon, is therefore likely to be incorrect, failing to generate the observed excess of 2Q-WPBs when modeled (**Figure 1D and 1E**). Alternatively, the ribbon might be formed from a collection of shorter, independent Golgi segments, mini-stack pairs included, which could act as preferential sites of formation for 2Q-WPBs. If *p* is sufficiently large, more 2Q-WPBs than 1Q-WPBs will form from paired mini-stacks; whereas at the lowest values of *p* only 1Q-WPBs will form. We tested whether a ribbon arrangement formed of short segments might explain the observed frequency of 2Q-WPBs.

We modeled a Golgi ribbon formed by elements of *l* = 3 (shown schematically in **Figure 2A**) and *l* = 4, which give rise to WPBs of size 1Q-3Q and 1Q-4Q, respectively, with the frequencies shown in **Table 1.**

**Figure 2:**
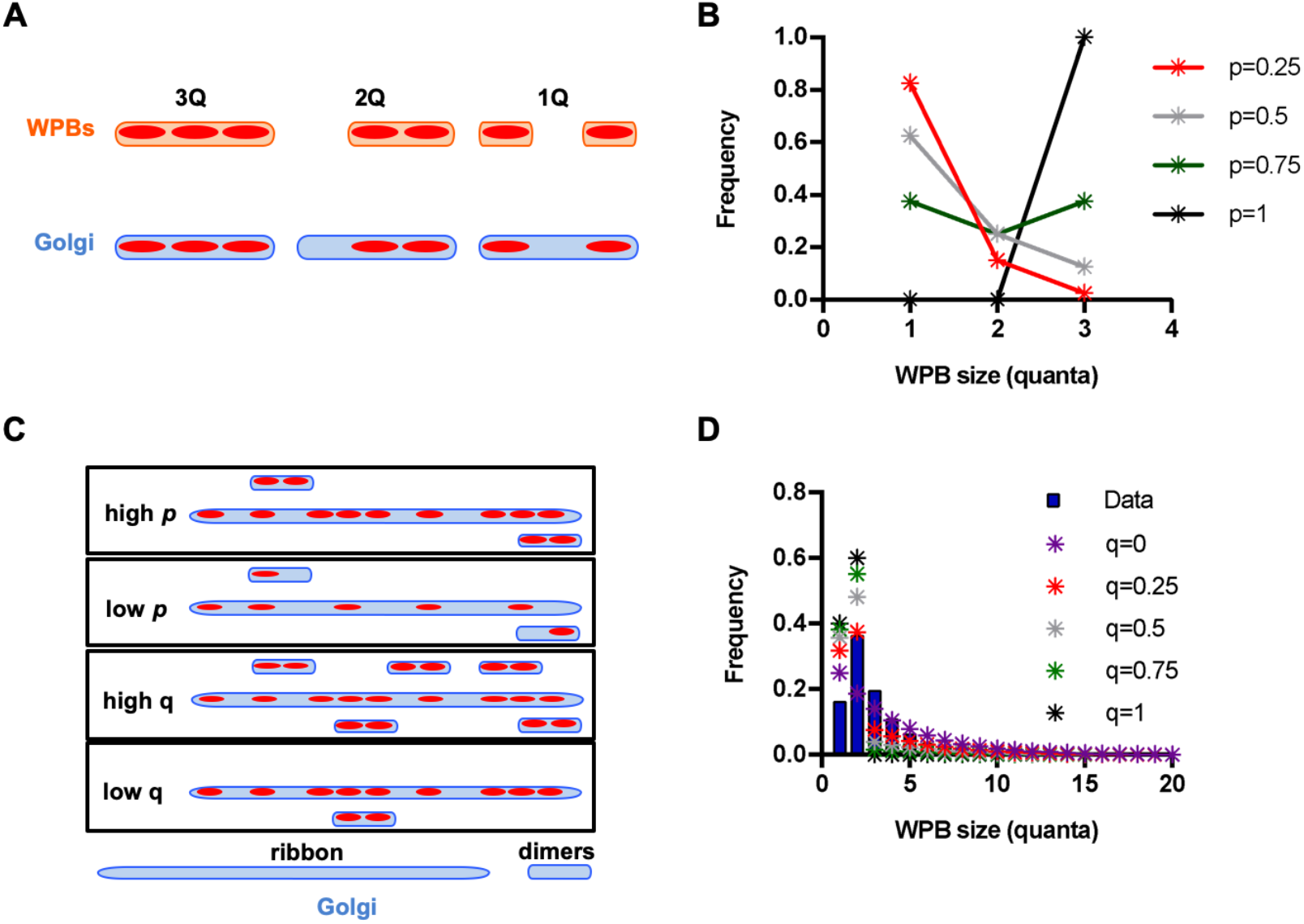
“Mini-ribbons” or “floating mini-stack dimers plus a long ribbon” models do not account for WPB size distribution. (A) Cartoon illustrating a model whereby the Golgi is composed of short ribbons of length 3. WPBs of size 1Q, 2Q or 3Q can be made depending on probability of mini-stack occupancy, *p*. (B) For a Golgi made as described in (A), the expected frequency distributions of 1Q-, 2Q- and 3Q-WPBs at the indicated values of *p* are shown. (C) Cartoon illustrating a model whereby the Golgi structure is made of long ribbons and mini-stack dimers, present in proportion *q*. (D) based on the model in (C) WPB size distributions calculated for *p* = 0.75 and the indicated values of *q* are shown (stars) in comparison to the measured size distribution (blue bars).

**Table 1.** Expected WPB size frequencies for ribbons formed by 3 and 4 ministacks linearly arranged. Modeling of WPB proportions generated by Golgi mini-ribbons made by 3 or 4 mini-stacks. Numerical solutions for indicated values of *p* are shown in Figure 2B and S2A.

|  | Proportion of<br>WPBs of<br>length 1 | Proportion of<br>WPBs of<br>length 2 | Proportion of<br>WPBs of<br>length 3 | Proportion of<br>WPBs of<br>length 4 |
| --- | --- | --- | --- | --- |
| Golgi length<br>$l = 3$ | $3p(1 - p)^2 + 2p^2(1 - p)$ | $2p^2(1 - p)$ | $p^3$ | 0 |
| Golgi length<br>$l = 4$ | $4(1 - p)^3p + 6p^2(1 - p)^2 + 2p^3(1 - p)$ | $3p^2(1 - p)^2 + 2p^3(1 - p)$ | $2p^3(1 - p)$ | $p^4$ |

In the first case, the maximum proportion of 2Q-WPBs predicted is 0.268 (for *p* = 0.75, **Figure 2B**), while in the second it is always < 0.3 (**Figure S2A**). Therefore, a Golgi “ribbon” arranged in segments of linked but independent mini-stack trimers and tetramers does not explain the observed 2Q-WPB frequency. Moreover, these scenarios predict once again that the frequency of 1Q-WPBs should exceed that of 2Q-WPBs at any value of *p*, except 1 (**Figure 2B and S2A**), and thus do not correctly simulate the experimental data.

From these simple modeling exercises it appears that the only way to get a proportion of 2Q-WPBs exceeding 0.3 is to have a significant number of functionally paired mini-stacks. Ultimately, if *p* = 1, then having a Golgi ribbon composed of mini-stacks linked into functional units whose length distribution is equal to the WPB length distribution will obviously match the data. However, the earlier mentioned cell-to-cell variability in VWF expression implies that at the cell population level the probability of mini-stack occupancy, *p*, cannot be 1.

Let us then assume that the Golgi ribbon consists of a mixture of mini-stacks arranged into functional dimers and long ribbon segments (depicted in **Figure S2B and Figure 2C**).

Equation **3** describes this scenario, where *q* represents the proportion of ministacks existing as dimers.

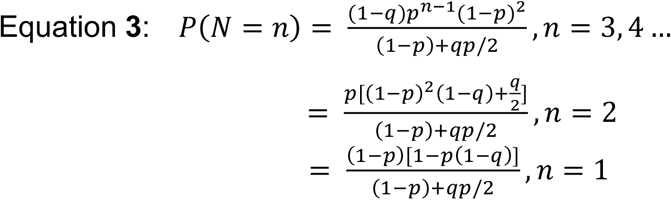

In conditions where occupancy probability, *p*, is > 0.5 and *q* is sufficiently high, this model does predict an excess of 2Q-WPBs compared to 1Q-WPBs (**Figure 2D** and **Figure S2C**). Nevertheless, in these scenarios the proportion of 3Q-WPBs is lower than that experimentally measured (**Figure 1C**). Thus, while the presence of separate mini-stack dimers and a long ribbon might account for the high proportion of 2Q-WPBs in the experimental data, it does not account for proportions of 1Q- and 3Q-WPB.

### 1Q-WPB instability

A major assumption built into our modeling so far is that WPBs of different sizes behave similarly; meaning that they are generated, stored and disappear from cells with identical kinetics. However, the unexpected frequency of 1Q-WPBs might be due to their being less stable than larger WPBs. We modeled this scenario by assuming that for every currently forming WPB of length greater than one, there are *β* times as many WPBs of that length in the cell, whereas for 1Q-WPBs, there are *β’* times as many. We can therefore derive a value, *α*:

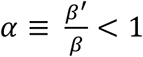

where *α* describes the relative instability of 1Q-WPBs. In this scenario, the proportion of 1Q-WPB is given by:

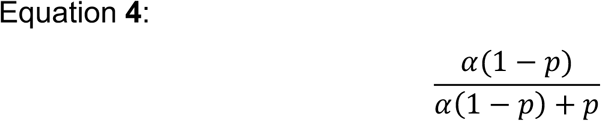

Whilst the proportion of WPB of length *N*, where *N* > 1, is:

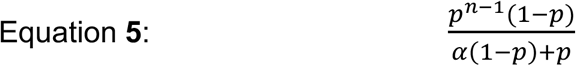

We can fit the value of *p* for this distribution to the experimental data by minimizing the Kolmogorov-Smirnoff (KS) distance, that is the difference between the theoretical and experimental distributions for *N*≥*2*. We find a best-fit value of *p* of 0.5692.

Using equation 4, we can then fix *α* to give us the correct proportion of 1Q-WPBs:

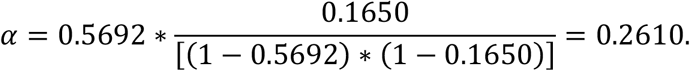

For these values of *p* and *α*, the model-predicted and the experimental distributions do indeed show excellent agreement (**Figure 3A**).

**Figure 3.**
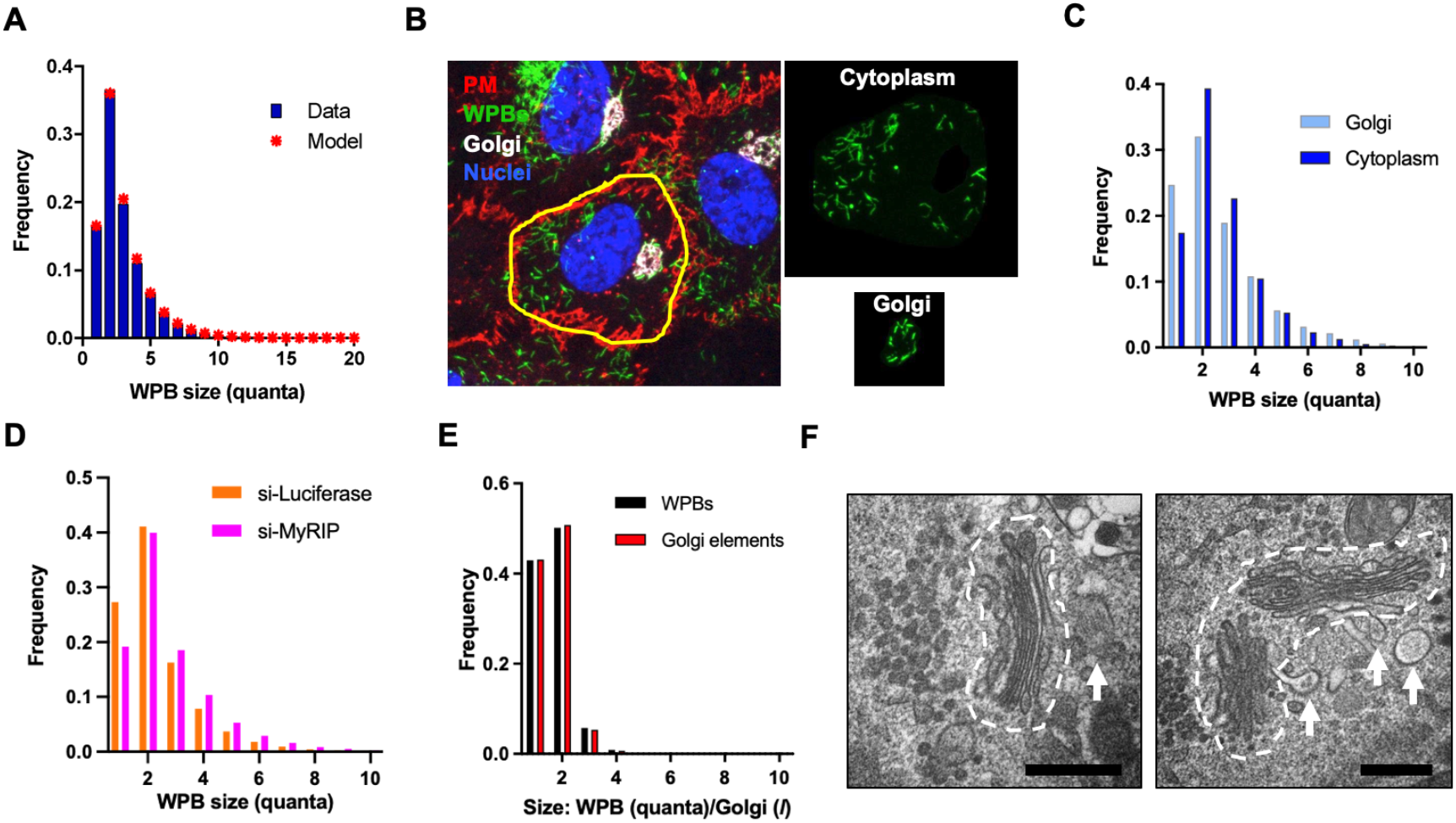
Instability of 1Q-WPBs and existence of stable mini-stack dimers. (A) Predicted WPB size distribution (stars) compared to real data (blue bars), where 1Q-WPBs display instability, α, greater than longer organelles. (B-C) HUVECs were fixed and processed to label the Golgi apparatus with an anti-TGN46 antibody and WPBs and cell boundaries as described in Figure 1B. The Golgi area was masked to identify newly-made WPBs and separate them from the cytoplasmic organelles (panels on the right). (C) Measured size distribution of Golgi and cytoplasm WPBs. (D) Size distribution of WPBs in Luciferase- or MyRIP-depleted cells. (E) After secretory granule depletion by PMA, cells were allowed to replete their WPB pool in nocodazole. Size distributions of Golgi elements and WPB were measured. (F) Electron micrographs of HUVECs treated with nocodazole for 24 h. Golgi mini-stacks visible as separate entities (dashed outline) were observed as monomers (left) or dimers (right); arrows point at forming WPBs. Bars: 500 nm.

These results prompted us to further examine the possibility of 1Q-WPBs’ instability. If 1Q-WPBs are preferentially lost after biogenesis, they might be present in higher proportions where they are assembled; i.e. near and at the Golgi. We imaged and separately quantified the size of WPBs present in the vicinity of the Golgi versus in the rest of the cytoplasm. The peri-Golgi pool of WPBs indeed shows a higher fraction of 1Q-WPBs compared to those imaged in the cell periphery (**Figure 3B** and **3C**). These data raise the question of potential mechanisms underlying the differential depletion of the 1Q-WPBs. Endothelial cells control which sizes of WPBs are selected for exocytosis (34). Therefore, one possibility is that 1Q-WPBs disappear by exocytosis. Release of VWF stored in WPBs occurs either acutely following exposure to agonist (stimulated exocytosis) or tonically, in the absence of any stimulus (basal exocytosis) (35). WPBs are anchored to actin fibers adjacent to the cell membrane via the effectors Rab27 and MyRIP. Ablation of either of these proteins increases the release of VWF through basal exocytosis (36). Does basal release differentially affect WPBs of different size classes? Depleting cells of MyRIP increases the fraction of long WPBs (**Figure S3A** and **S3B**), changing WPB size distribution (**Figure 3D).** The reduced proportions of 1Q-WPBs indicate that secretory granules of this size class are preferentially depleted by basal exocytosis, and thus more readily disappear from the cell than longer organelles. The relative peri-Golgi enrichment of 1Q-WPBs and their differential propensity to undergo basal exocytosis thus, at least partly, explain the observed secretory granule size distribution.

### Stable mini-stack dimers

Our WPB biogenesis model posits that organelle length is determined by VWF quantum occupancy of adjacent mini-stacks (statement *iii*). Microtubules, their regulators and dynein motors are necessary for mini-stack tethering and Golgi ribbon formation (18, 22, 23). Nocodazole depolymerizes microtubules, leading to ribbon unlinking into mini-stacks (22, 37). We thus set out to quantify the effects of nocodazole treatment on WPB size distribution. Exposure to the strong secretagogue PMA (phorbol 12-myristate, 13-acetate) clears cells of pre-existing WPBs (**Figure S3C**) (38). We implemented this treatment followed by recovery in the presence of nocodazole for 24h (**Figure S3D**). Compared to that of DMSO-pretreated and nocodazole-chased cells, the WPB size distribution of PMA-pretreated and nocodazole-chased cells shows a dramatic fall in the WPB classes > 2Q, compensated by a marked increase in 1Q-WPBs; remarkably, the proportion of 2Q-WPBs was unchanged (**Figure S3E**). Golgi fragments generated by nocodazole treatment are scattered throughout the cell (**Figure S3D**), facilitating their morphometric quantification. Golgi elements were thus measured and rounded up to the nearest multiple of *l*, as done for WPBs. This analysis revealed that Golgi elements were almost exclusively of sizes 1*l* and 2*l*, closely matching the distribution of WPB size classes (**Figure 3E**). These data are consistent with repeated observations that the Golgi ribbon is necessary to the formation of long WPBs (29, 32, 39); most importantly, they strongly suggest that even in the absence of microtubules, the vertebrate Golgi is not reduced exclusively to individual mini-stacks, but that around 50% of the Golgi objects generated by nocodazole treatment are size consistent with mini-stack dimers.

Inspection of the ultrastructure of the Golgi apparatus indeed showed the frequent occurrence of paired mini-stacks following nocodazole treatment, indicating they are stable structures in the absence of microtubules (**Figure 3F and S3F**).

### The ribbon as a mixture of mini-stack monomers and dimers

Based on findings above, we tried to assess how strong the preference for the mini-stack dimer vs monomer configurations might be by modeling binding rates of Golgi elements. We assumed that in the presence of intact microtubules, the Golgi elements continuously undergo a linking/unlinking process; that the rate at which two elements join is independent of their size; and that unlinking is slower for mini-stack dimers.

Let the binding rate be *r*, and the unbinding rate *d_1_* if it does not split a stable dimer, and *d_2_* < *d_1_* if it does. To simplify the modeling, we assume that two neighboring monomers always form a dimer, so that in longer sections of the ribbon, dimers can be separated, at most, by one monomer. We note that tetramers can have two forms either consisting of two coupled dimers or a dimer with a monomer on either side. To make the system of equations finite, we also ignore portions of ribbon with lengths > five. We let the number of portions of length n be *l_n_*, for n = 1 to 5, and let N be the total number of mini-stacks. We denote by *l_4_*, the number of tetramers consisting of two dimers and *l_4_’* the number of tetramers consisting of a dimer and two monomers. There are two types of pentamer: each has two dimers and one monomer, but the monomer can be on the end or in the middle. These need to be considered separately, since they have different unbinding rates to the two types of tetramer (**Figure S4**). We denote the number of pentamers with a monomer on the end by *l_5_* and the number with a monomer in the middle by *l’_5_*. We obtain the reaction system given in **Figure S5A**. We can also represent the evolution of the mean numbers of each type of Golgi piece by differential equations, see equations **6** (**Figure S5B**). The final values of the numbers of Golgi fragments of different lengths are given by the steady state values of these equations. Depending on the values of the parameters *r*, *d_1_*, *d_2_* and the total number of mini-stacks, the distributions can peak at dimers and can have trimers either more or less frequent than tetramers. This is illustrated by stochastic simulations of the system of reactions, changing the value of the unbinding rate d (Matlab code provided in **Figure S6**). Two examples are shown: in **Figure 4A**, pieces of length three are more abundant than those of length four in one simulation, whereas in the other the opposite is true.

**Figure 4.**
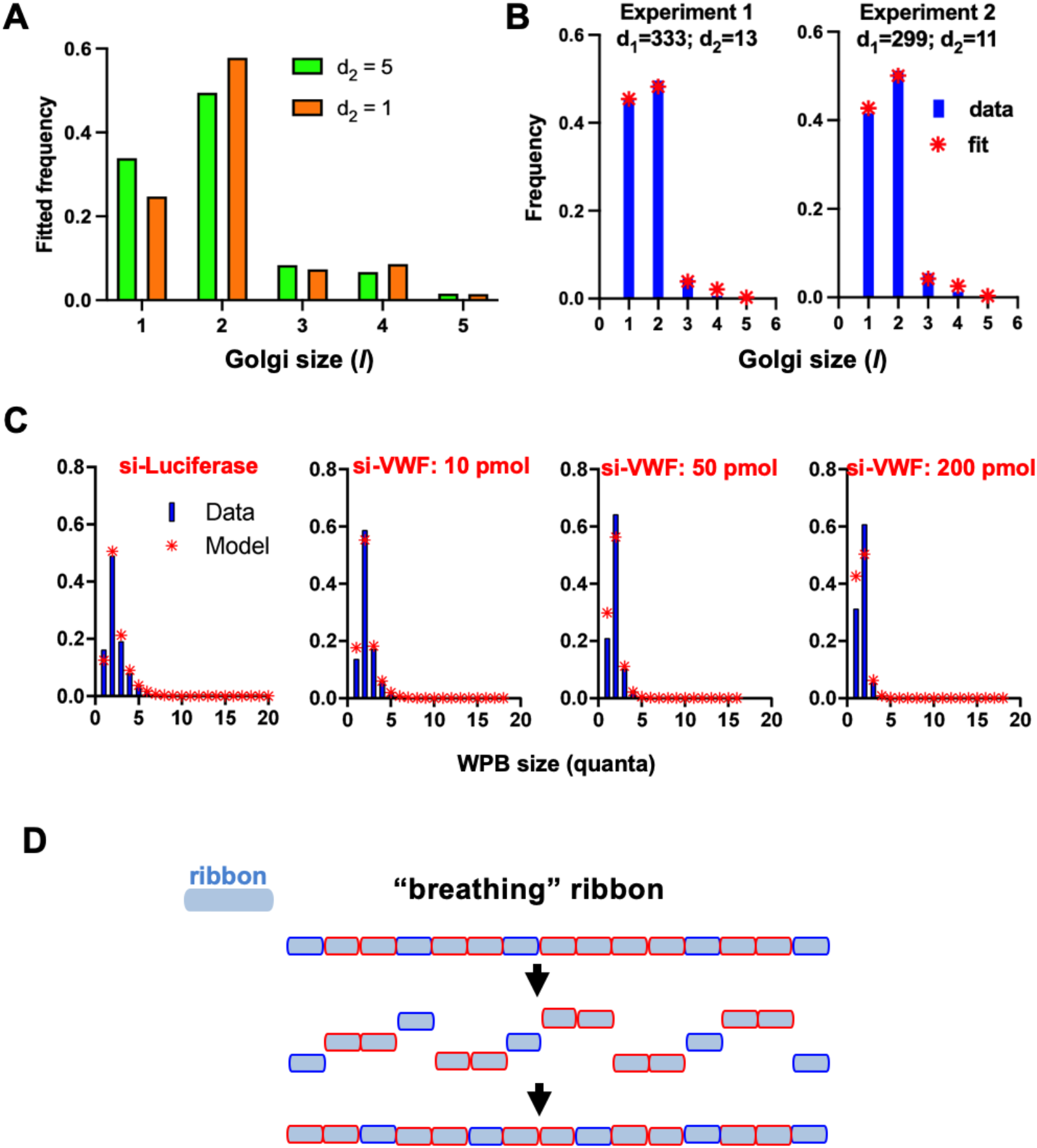
A breathing model of the Golgi ribbon. (A) Modeling mini-stack binding and unbinding rates generated by solving an equations system based on a reaction scheme (Figure S5). Two examples of fitted frequencies of Golgi of size 1-5 *l* (i.e., number of mini-stacks) obtained by posing r = 0.001, d_1_ = 100, total number of mini-stacks = 100000 and d_*2*_ at the indicated values. (B) Relative frequencies of Golgi pieces of different lengths (in mini-stacks) from two experiments with nocodazole-treated cells, together with the predictions from the model with the best fit values of d_1_ and d_2_ (each fitted to the nearest whole number). (C) Predicted length distribution of WPBs (red stars), maintaining α constant between datasets, compared to real data (blue bars) in control (si-Luciferase) and different levels of VWF expressing cells (si-VWF). Note: the higher the amount of siRNA to VWF, the lower its expression. (D) The “breathing” Golgi ribbon model is schematized. Occurrence of weak bonds between monomers and stable dimers predicts their dynamic linking/unlinking and likely rearrangement within the ribbon. The model shows that the number of stable mini-stack dimers is the same as that of monomers as measured in our experiments (Figure 3E-EF).

Whether tighter linking of dimers leads to an abundance of Golgi pieces of length four (and potentially all multiples of two) therefore depends on the parameters of the system. We now fit the data. We are interested in the steady state frequencies of Golgi pieces of different length; therefore, in the model, we are free to choose both the total number of Golgi pieces (although this should be large to accurately sample the predicted frequencies) and the rate at which the system reaches equilibrium. The only combinations of parameters that should affect the steady state frequencies are rN/d_1_ and d_1_/d_2_. We therefore only need to explore different values of these two dimensionless parameter combinations. We can do so by arbitrarily fixing N (subject to it being large enough to get good sampling) and *r* and varying *d_1_* and *d_2_*. Therefore we fix *r* = 0.001 and N = 100,000 (modeling a population of cells), both of these are dimensionless, and fit *d_1_* and *d_2_* to the data on proportions of Golgi of different lengths. We show in **Figure 4B** the distributions of lengths of Golgi fragments from two actual experiments, in which cells were treated with nocodazole, and from the model fit to those experiments. The reason we used these data is that in the control conditions, i.e. in the context of a linked ribbon, individual Golgi elements cannot be accurately measured, while this is possible after nocodazole treatment (**Figure 3E** and **S3D**). These simulations show a good fit to the experimental data. The calculated values of the dissociation rates based on the experimental data are *d_1_* = 333 and 299 and *d_2_* = 13 and 11 (chosen to minimize KS distance), respectively. Thus the prevalence of dimers may be explained by a striking 30-fold difference in mini-stack dissociation rates. We note that the actual values of *d_1_* and *d_2_* depend on our choice of the dimensional time corresponding to a unit of dimensionless time, which we cannot estimate from the steady state proportions of Golgi lengths.

Having approximately 50% of the Golgi ribbon composed of (possibly 30-fold) more-tightly linked dimers might explain why nocodazole fails to unlink the Golgi entirely into monomers (**Figure 3E-3F** and schematized in **S3F**). Unlike an earlier model (**Figure 2C-2D and S2C-S2D**), where we considered ministack dimers as existing separate from the ribbon, by having 50% of Golgi objects as functional dimeric mini-stacks that are incorporated into the ribbon (**Figure S4)**, it is possible to assemble longer organelles on these. If this is the case, then according to our model, lowering the number of quanta within the system without changing ribbon structure should reveal the number of larger WPBs diminishing as the general chances of co-packaging fall whilst the number of 2Q-WPBs should be less affected. The lowered chances of copackaging will also increase the fraction of 1Q-WPB, assuming that their instability is unaffected. Endothelial cells were treated with different amounts of VWF-targeting siRNA to reduce its cellular levels and therefore the number of quanta, without changing quantum size (29), therefore lowering *p* (**Figure 4C**). We note that the model fits are less good than in **Figure 3A**, because we have required α to be constant across all four conditions, where it thus has to explain more data. Allowing α to vary between the conditions results in much better fits (data not shown; we note that the best fit value of α decreased with increasing siRNA concentration), but we could not justify this assumption biologically. We find that even when VWF levels are at their lowest (**Figure 4C**, si-VWF: 200 pmol siRNA), the proportion of 2Q-WPBs remains high, suggesting that dimers are a preferred unit of assembly.

These results suggest that the excess of 2Q-WPB does reflect the presence of functionally different mini-stacks dimers within the ribbon. From this follows that the assumption that mini-stack linkages are identical and thus that quantal co-assembly in WPB formation is a purely random process (assumption **c**) is not likely to be true.

## Discussion

Weibel-Palade bodies represent a unique cellular system due to the two-tiered functioning of the Golgi apparatus in their biogenesis. Golgi mini-stack dimensions constrain the size of VWF quanta, while the Golgi ribbon architecture controls co-packaging of VWF quanta into nascent secretory granules (29). As a consequence, the structural arrangement of a biosynthetic compartment, the Golgi, is closely reflected by the size of the organelles it generates, the WPBs. In other words, the size distribution of WPBs provides a proxy for the architecture of the Golgi apparatus. Taking advantage of this unique cellular system we generated a formal description of the structural organization of the Golgi apparatus. Hypotheses about the Golgi ribbon structure, translated into mathematical modeling, predicted WPB size distributions that were tested against those actually measured using an image analysis workflow that we previously employed in a series of studies (29, 32, 34, 39–42).

The measured size distribution of WPBs posed two challenges to modeling of the Golgi ribbon structure. First, why are 1Q-WPB depleted with respect to other size classes? Second, what structure can explain the high frequency of 2Q-WPBs with respect to longer size classes?

For the Golgi ribbon structure to be consistent with both these data, and what is known of WPB biogenesis, the model had to be refined from that of a single very long ribbon made by linearly linked mini-stacks, through to that of an ensemble of short ribbon stretches, and finally, to an organization involving a mixture of mini-stack monomers and dimers within the ribbon. As for 1Q-WPBs, mathematical modeling suggested their low frequency might be explained by a selectively shorter residence time. Experimental testing of this hypothesis indeed indicates that 1Q-WPBs undergo basal exocytosis at higher rates than longer WPBs. In agreement with this finding, we had previously found that short WPBs are more prone to exocytosis (29, 32); the modeling exercise deployed in this study allowed us to narrow down this effect to 1Q-WPBs, as those preferentially selected for basal secretion.

Regarding the second question, the observed steady-state frequency of 2Q-WPBs can, in principle, be explained by assuming that the Golgi ribbon contains a large proportion of mini-stack dimers. This prompted experimental testing combining treatment with PMA and nocodazole, which deplete the cellular pool of WPBs and break up the ribbon into mini-stacks, respectively. Strikingly, in these conditions we observed that 1Q- and 2Q-WPBs were almost exclusively made, with their size matched by that of Golgi elements. Since there is no evidence from the literature for quantization of mini-stack size, we had to conclude that microtubule disruption generates Golgi elements existing as mini-stack monomers and dimers. This conclusion was confirmed by electron microscopy observations. Thus, a structural basis must exist to prevent mini-stack dimer dissociation. Based on the ratio of mini-stack monomers and dimers following nocodazole, we run simulations that resulted in an estimated “strength” of the link between mini-stacks in a dimer being approximately 30-fold greater than that of monomers with other monomers or dimers in the context of a ribbon (**Figure S4**, model). The modeling of our experimental data thus fits with a picture of the Golgi ribbon as a dynamic structure formed by mini-stack monomers and very stable dimers, accounting for the measured frequency of 2Q-WPBs.

Intriguingly, actin-dependent paired mini-stacks were described in insect cells, which display a Golgi organization typical of invertebrate animals, with scattered elements. These authors also showed that in mammalian HeLa cells, pretreated with nocodazole to unlink the ribbon, inhibition of actin polymerization caused Golgi elements’ scission, therefore surmising the presence of mini-stack pairs in mammalian cells (43). Our modeling exercise and the experiments it prompted indeed support the presence of stable, microtubule-independent mini-stack dimers in mammalian cells. At this stage it is not clear how these dimers are kept together. Based on the above-mentioned work (43), the actin cytoskeleton is the likely driver of mini-stack dimer formation. However, which of its regulators are involved remains to be established. Golgi membranes certainly provide a scaffold for many candidates (44).

Adopting WPB size distribution as a proxy for modeling the Golgi ribbon organization thus paints an unanticipated picture of this architecture. Our best-fitting model is consistent with a ribbon made from similar numbers of ministack monomers and dimers undergoing continuous linking/unlinking (**Figure 4D**). This “breathing” would make the Golgi ribbon a highly dynamic structure, capable of promptly responding to signals that require its rearrangement, such as re-positioning and re-orientation during migration and directed secretion (23, 45, 46). It is possible that such a structural organization may not be universal to vertebrate cells, but rather reflect an endothelial adaptation of the Golgi ribbon to the need for WPB production, providing a flexible system capable of modulating the size of these organelles in response to environmental and pharmacological cues (29, 34, 39, 41). However, the above-mentioned evidence of similar dimers present in HeLa cells (43) suggests that this is a more widespread feature. In our model, formation of a dimer prevents a similar strong interaction of either of the constituent mini-stacks with another monomer. This implies that the machinery responsible for stable dimer formation must segregate at the location of the mini-stacks interaction (**Figure S4)**; that is at the rims of the cisternae. Polarization of molecular machinery within organelles has been described. For instance, MyRIP, which interestingly interacts with Factin, localizes at one tip of WPBs (36). In mini-stack monomers, Golgi structural proteins localize at the rims of the cisternae (47) and one possibility is that lipid-mediated phase separation and consequent protein spatial restriction (48) may segregate the machinery involved in dimer stabilization.

Our modeling-based prediction of functionally distinct and stable mini-stack dimers does not shed light on the physical nature of their embedding into the ribbon superstructure. Fluorescence recovery after photo-bleaching (FRAP) experiments firmly established that cisternal membranes of neighboring mini-stacks are continuous (16, 17, 49). Therefore, a question raised by the model of the Golgi ribbon structure here described is what molecular or physicochemical characteristics locally define the mini-stack dimers, while allowing their membrane continuity with adjacent mini-stacks. Our findings may prompt further studies to investigate this issue.

In summary, our modeling exercise provides insight into the dynamics of the smaller class of WPBs. Further detailed analysis - using the quantitative model that we introduce in this study - of the roles of the full range of identified endothelial exocytic machinery would identify the molecular mechanisms regulating basal secretion and thus just how the 1Q-WPB are preferentially released, potentially critical to setting the level of basally secreted plasma VWF and thus thresholds for hemostasis (35). Perhaps of more general interest and importance, if our model were to reflect an actual structural arrangement, a systematic approach designed to identify the machinery controlling the architecture of the Golgi ribbon, using WPB size distribution as proxy of the mini-stack linkage status, seems likely not only to further our understanding of how this remarkable structure is formed, but also what might link its fragmentation to disease.

## Materials and Methods

### Cells, reagents and treatments

Human umbilical vein endothelial cells (HUVECs) were obtained commercially from PromoCell or Lonza. Cells were expanded and maintained as described previously (29). Chemicals were from Sigma-Aldrich. Cells seeded in 96-well plates were subjected to the described treatments, fixed and subjected to immuno-staining, followed by image acquisition and processing to extract organelle size through an analytical pipeline described in (29); script available on request. The size in μm of ~ 100000 WPBs was measured for each condition. Sample processing for electron microscopy was essentially as in (50). Procedures, antibodies and siRNA used are detailed in the SI appendix.

### Mathematical modeling

To create the model fit for Figure 3A, we used equation 5. We initially ignored the frequency of 1Q-WPBs. The relative frequencies of WPBs of lengths 2Q, 3Q, 4Q… are then given by 1-p, p(1-p),p^2^(1-p), … For multiple values of p in the range [0,1], we generated this distribution and measured the Kolmogorov-Smirnoff distance (this is the maximal absolute difference between the cumulative distribution functions) to the experimental relative frequency distribution of WPBs of lengths 2Q, 3Q, 4Q… We found the value of p that minimised this distance. Then, using equation 4 with the optimal value of p already found, we fixed α to give us the exactly correct frequency of 1Q-WPBs. To create the model distributions in Figures 4A and B, we fixed values for *d_1_* and *d_2_* and then ran a code to numerically simulate the stochastic process described by the reactions given in Figure S5 (see Figure S6 for the code). At a late time (t=5000), we output the relative frequencies of the Golgi portions of length 1-5 mini-stacks. For Figure 4B, we then computed the Kolmogorov-Smirnoff distance to the experimental values of those relative frequencies. We repeated the process for *d_1_*=1, 2, 3, …, 20 and *d_2_*=1, 2, …, 400 and found the values of *d_1_* and *d_2_* that minimized the distance. Note that had these values been at an extreme value of the range, we would have searched a broader set of values for *d_1_* and/or *d_2_*.

To create the model fits for Figure 4C, we fit values of p to the data from the control and three siRNA concentration experiments, as we did for Figure 3A. We then used the frequencies of 1Q-WPBs to exactly determine alpha for each experiment. However, we assumed that α would not vary due to siRNA treatment, so that the values of α should be the same for all conditions. Thus, we took the weighted mean of the values of α that best fit each data set. We multiplied the value of α obtained for the luciferase siRNA condition by the total number of WPBs in that experiment and added the values of α obtained for VWF siRNA at 10-200 pmol, multiplied by the corresponding number of WPBs in each condition, then divided by the total number of WPBs over all four conditions. We then plotted the model predicted distributions with that single value of α and the values of p chosen for each experiment separately. Note that had we allowed α to vary between the conditions, the fits would have been considerably better.

## Supporting Information

### Detailed Materials and Methods

#### Cells

Human umbilical vein endothelial cells (HUVECs) pooled from donors of both sexes were obtained commercially from PromoCell or Lonza. Cells were expanded and used within 15 population doublings, corresponding to passage 3 to 4 after expansion by our lab and maintained in HUVEC Growth Medium (HGM): M199 (Gibco, Life Technologies), 20% Fetal Bovine Serum, (Labtech), 30 μg/mL endothelial cell growth supplement from bovine neural tissue and 10 U/mL Heparin (both from Sigma-Aldrich). Cells were cultured at 37 °C, 5% CO_2_, in humidified incubators

#### Reagents

Chemicals. Nocodazole (M1404), Phorbol 12-myristate 13-acetate (P8139) and DMSO (D2650) were from Sigma-Aldrich. Nocodazole and PMA were dissolved in DMSO to generate 10 and 0.1 mg/ml stock solutions, respectively. Antibodies used in this study were: rabbit polyclonal anti-VWF propeptide region (1); mouse monoclonal anti-VE-cadherin (BD Bioscience, Clone 55-7H1, cat. no. 555661); sheep anti-TGN46 (BioRad, cat. No, AHP500G). siRNAs targeting firefly Luciferase (5’-CGUACGCGGAAUACUUCGAdTdT-3’), human VWF (5’-GGGCUCGAGUGUACCAAAAdTdT-3’) and human MyRIP (5’-AAGGTGGGAATTATTATTTAA-3’ and 5’-CCAAATTTACTTCCCAATAAA-3’) were previously described and validated (27, 34). siRNAs were custom synthesized by Eurofins MWG Operon.

#### Treatments

HUVECs were seeded on gelatin-coated NUNC 96-well plates (Thermo Fisher, cat.no. 167008) at 15000-20000 cells/well and maintained in HGM. Untreated cells were fixed after 48 h. In WPB repopulation experiments, 23 h after seeding, cells were fed with HGM supplemented with either DMSO or with PMA (100 ng/ml) for 1 h to elicit massive VWF secretion and deplete WPBs, followed by nocodazole incubation (1 μg/ml) for 24 h to allow WPB formation from separated Golgi mini-stacks. In the experiments involving VWF or MyRIP depletion, the indicated amounts of siRNAs were electroporated into one million HUVECs per reaction with an AMAXA Nucleofector II (Lonza). Cells were then plated in 96-well plates as described above and cultured for 48 h. Cells were then rinsed twice with warm, fresh HGM and fixed by incubation with 4% formaldehyde in PBS (10 min, at room temperature).

#### Immuno-staining and image acquisition

After permeabilization (with 0.2% TX-100 in PBS) and blocking (with 1 % bovine serum albumin in PBS), cells were incubated with primary antibodies, followed by secondary AlexaFluor dye-conjugated antibodies (Life Technologies), both diluted in 1% BSA, 0.02% TX-100 in PBS. Nuclei were counterstained with Hoechst 33342 (Life Technologies, H3570). Imaging was carried out with an Opera High Content Screening System (Perkin Elmer), using a 40x air objective (NA 0.6). Nine fields of view per well in 8-16 replicate wells were acquired, corresponding to approximately 2400-4800 cells per condition. In some experiments, together with WPBs, the Golgi apparatus and cell boundaries were visualized with anti-TGN46 and VE-cadherin antibodies.

#### Image analysis

High-throughput morphometry (HTM), the analytical pipeline including image processing, organelle segmentation and measurement of morphological parameters, has been described in detail elsewhere (2). Briefly, images of WPB (visualized with anti-VWF propeptide antibody) were segmented using Python (v2.7) (script available upon request). Data analysis was done using *R* (i386 3.1.0). The size in μm of ~ 100000 WPBs was measured for each condition. WPB lengths were expressed as numbers of quanta as described in the Results section.

#### Electron microscopy

Nocodazole treated cells were high-pressure frozen, free-substituted (HPF/FS) and processed as described previously (3). Samples were imaged with FEI Tecnai 20 TEM using an Olympus-Soft Imaging System Morada camera.

**Figure S1.**
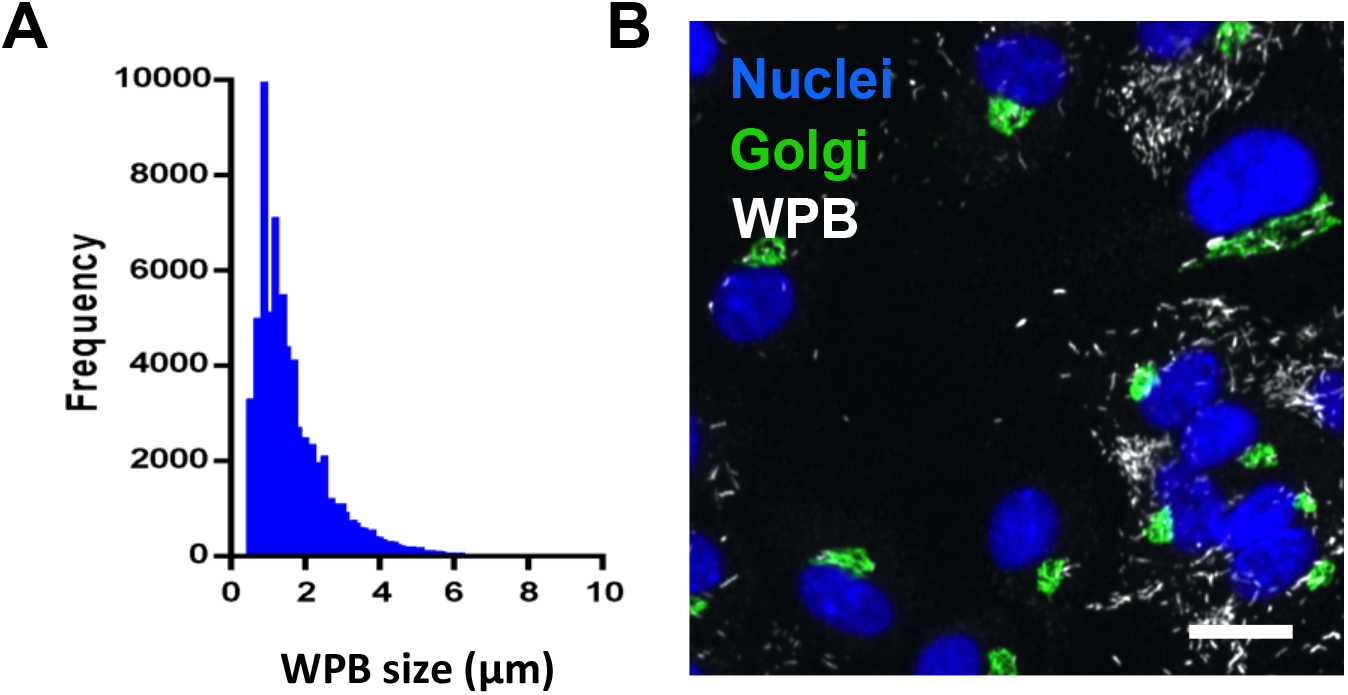
WPB size distribution measured from a population of endothelial cells. (A) WPB size distribution as measured by high-throughput morphometry (see SI Material and Methods). (B) HUVECs show variability in VWF expression and WPB production. WPBs and nuclei were visualized as indicated in Figure 1B; the Golgi was visualized with an anti-GM130 antibody; scale bar, 20 μm.

**Figure S2.**
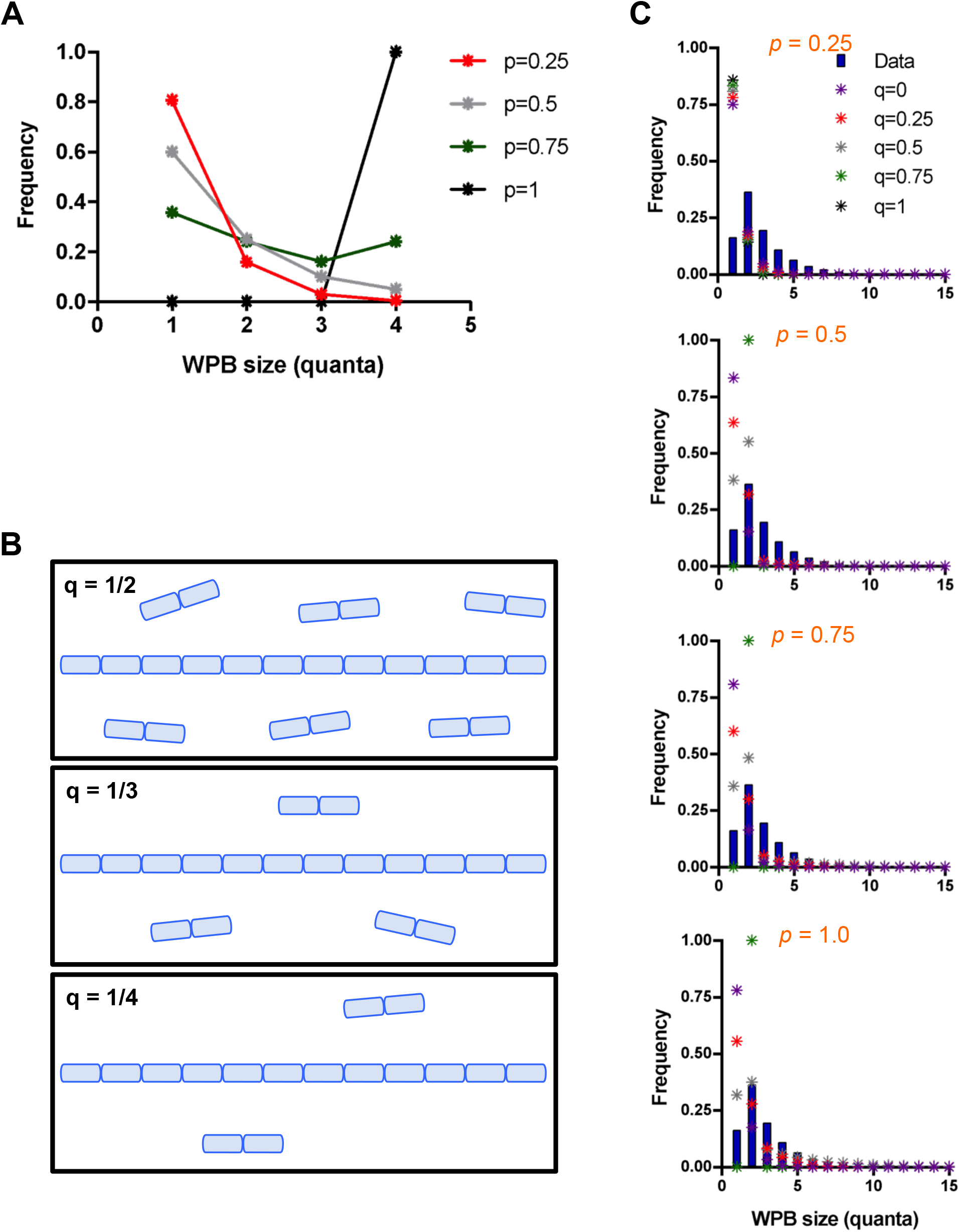
Further simulations based on the “mini-ribbons” and the “floating mini-stack dimers and long ribbon” models. (A) Predictions of WPB size distributions at different probabilities of quantum mini-stack occupancy probability (*p*), where the Golgi is made exclusively of ribbons of length 4*l*. (B) A Golgi model made of ribbons and mini-stack dimers, where the latter are present in proportion q; different q values are depicted. (C-E) Based on the model in B, expected WPB size distributions (stars) were calculated for the indicated mini-stack occupancy probability (*p*) and the proportion of the Golgi formed by dimers (q) and compared to the measured WPB size distribution (blue bars).

**Figure S3.**
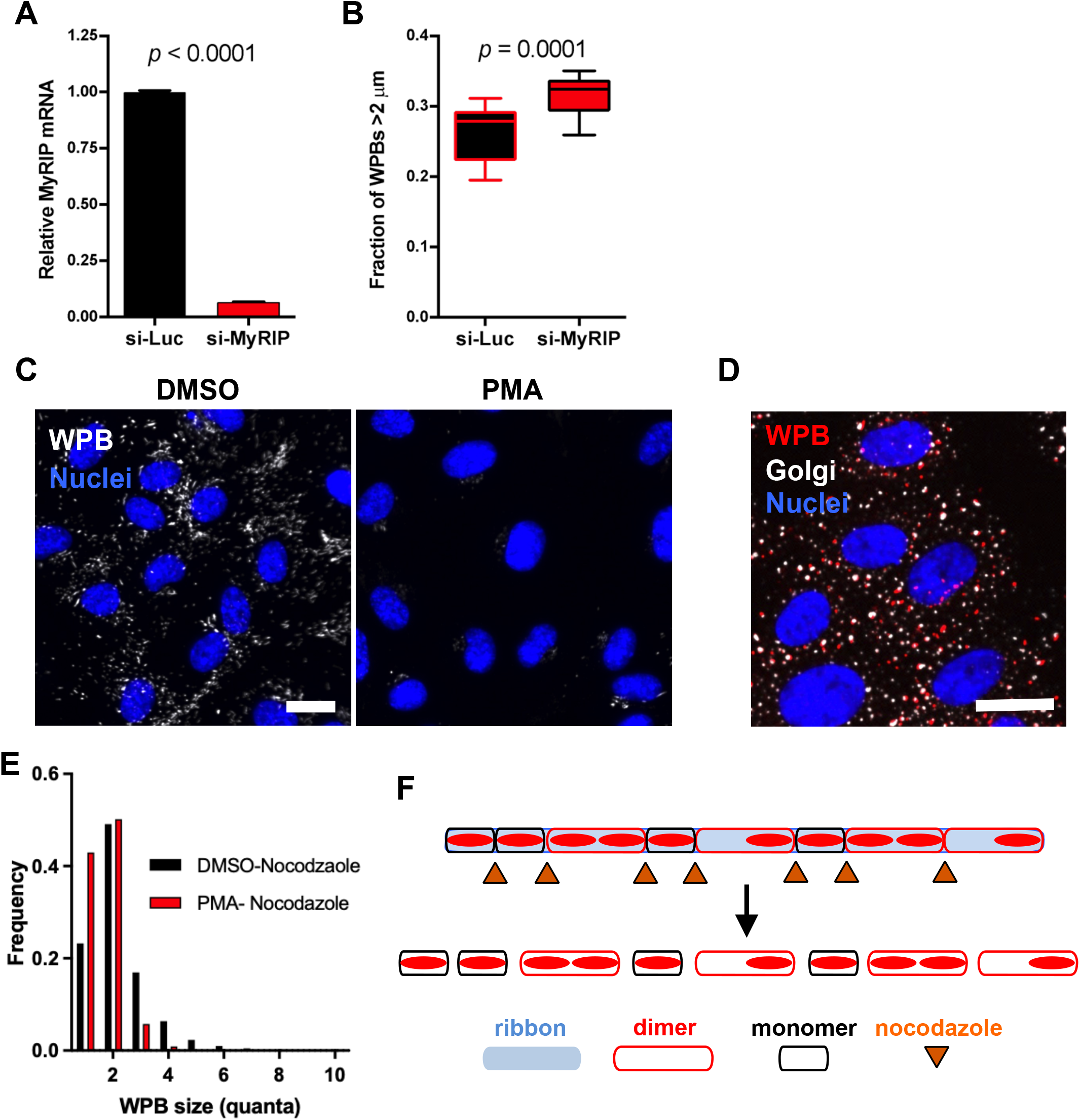
Shifts in WPB size distribution following increase in basal secretion and Golgi ribbon unlinking. (A) Efficiency of MyRIP knockdown (300 pmol of each siRNA were used per reaction). (B) Morphometric analysis of WPB size shows that MyRIP knockdown increases the fraction of long organelles (defined as those > 2 μm). (C) HUVECs were treated for 1 h with either DMSO or 100 ng/mL PMA and processed for immunofluorescence. PMA almost completely depletes the cellular pool of WPBs. Scale bar, 10 μm. (D) Representative micrograph of HUVECs pretreated with PMA as in (A) to clear WPBs and then chased in nocodazole for 24 h. Scale bar 20 μm. (E) Size distribution of WPBs 24h after nocodazole treatment, following pretreatment with either DMSO (control) or PMA to deplete the organelles. The resulting populations of organelles following 24h nocodazole treatment are: newly-made WPBs plus those left after basal exocytosis, in the case DMSO; almost completely newly-made WPBs, in the case of PMA. (F) Visualization of a hypothetical arrangement of mini-stacks within the Golgi ribbon, with the existence of stable mini-stacks dimers that are independent of microtubules.

**Figure S4.**
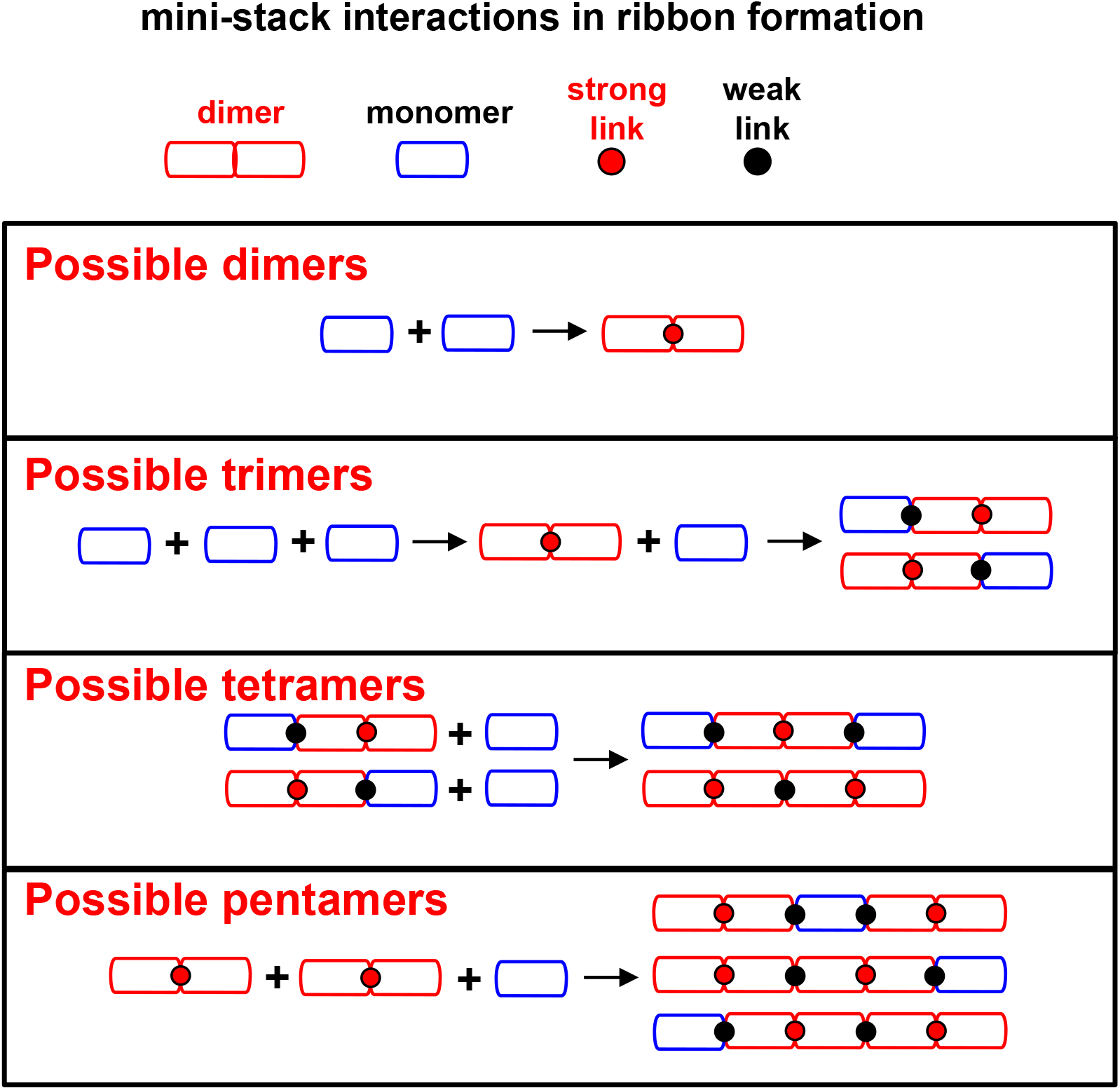
Modeling of mini-stack linking into the Golgi ribbon. Based on the modeling assumptions (see text), the possible mini-stacks associations up to pentamers are depicted.

## References

1. M. S. Vildanova, W. Wang, E. A. Smirnova, Specific organization of Golgi apparatus in plant cells. Biochemistry (Mosc) 79, 894–906 (2014).

2. H. Yano et al., Distinct functional units of the Golgi complex in Drosophila cells. Proceedings of the National Academy of Sciences of the United States of America 102, 13467–13472 (2005).

3. M. Sato et al., Caenorhabditis elegans SNAP-29 is required for organellar integrity of the endomembrane system and general exocytosis in intestinal epithelial cells. Molecular biology of the cell 22, 2579–2587 (2011).

4. N. Nakamura, J. H. Wei, J. Seemann, Modular organization of the mammalian Golgi apparatus. Curr Opin Cell Biol 24, 467–474 (2012).

5. J. H. Wei, J. Seemann, Unraveling the Golgi ribbon. Traffic (Copenhagen, Denmark) 11, 1391–1400 (2010).

6. J. H. Wei, J. Seemann, Golgi ribbon disassembly during mitosis, differentiation and disease progression. Curr Opin Cell Biol 47, 43–51 (2017).

7. D. A. Thayer, Y. N. Jan, L. Y. Jan, Increased neuronal activity fragments the Golgi complex. Proceedings of the National Academy of Sciences of the United States of America 110, 1482–1487 (2013).

8. S. Ireland et al., Cytosolic Ca(2+) Modulates Golgi Structure Through PKCalpha-Mediated GRASP55 Phosphorylation. iScience 23, 100952 (2020).

9. C. F. Daussy et al., The inflammasome components NLRP3 and ASC act in concert with IRGM to rearrange the Golgi during HCV infection. J Virol 10.1128/JVI.00826-20 (2020).

10. P. Gosavi, F. J. Houghton, P. J. McMillan, E. Hanssen, P. A. Gleeson, The Golgi ribbon in mammalian cells negatively regulates autophagy by modulating mTOR activity. Journal of cell science 131 (2018).

11. S. E. Farber-Katz et al., DNA damage triggers Golgi dispersal via DNA-PK and GOLPH3. Cell 156, 413–427 (2014).

12. S. Bellouze et al., Stathmin 1/2-triggered microtubule loss mediates Golgi fragmentation in mutant SOD1 motor neurons. Molecular neurodegeneration 11, 43 (2016).

13. C. Rabouille, G. Haase, Editorial: Golgi Pathology in Neurodegenerative Diseases. Frontiers in neuroscience 9, 489 (2015).

14. G. Joshi, Y. Chi, Z. Huang, Y. Wang, Abeta-induced Golgi fragmentation in Alzheimer’s disease enhances Abeta production. Proceedings of the National Academy of Sciences of the United States of America 111, E1230–1239 (2014).

15. P. Gosavi, P. A. Gleeson, The Function of the Golgi Ribbon Structure - An Enduring Mystery Unfolds! Bioessays 39 (2017).

16. T. N. Feinstein, A. D. Linstedt, GRASP55 regulates Golgi ribbon formation. Molecular biology of the cell 19, 2696–2707 (2008).

17. M. A. Puthenveedu, C. Bachert, S. Puri, F. Lanni, A. D. Linstedt, GM130 and GRASP65-dependent lateral cisternal fusion allows uniform Golgi-enzyme distribution. Nat Cell Biol 8, 238–248 (2006).

18. S. Yadav, M. A. Puthenveedu, A. D. Linstedt, Golgin160 recruits the dynein motor to position the Golgi apparatus. Developmental cell 23, 153–165 (2012).

19. Y. Zhang, J. Seemann, Rapid degradation of GRASP55 and GRASP65 reveals their immediate impact on the Golgi structure. The Journal of cell biology 220 (2021).

20. T. M. Witkos, M. Lowe, The Golgin Family of Coiled-Coil Tethering Proteins. Frontiers in cell and developmental biology 3, 86 (2015).

21. P. Kulkarni-Gosavi, C. Makhoul, P. A. Gleeson, Form and function of the Golgi apparatus: scaffolds, cytoskeleton and signalling. FEBS Lett 593, 2289–2305 (2019).

22. J. Thyberg, S. Moskalewski, Microtubules and the organization of the Golgi complex. Exp Cell Res 159, 1–16 (1985).

23. P. M. Miller et al., Golgi-derived CLASP-dependent microtubules control Golgi organization and polarized trafficking in motile cells. Nat Cell Biol 11, 1069–1080 (2009).

24. D. Tang et al., Mena-GRASP65 interaction couples actin polymerization to Golgi ribbon linking. Molecular biology of the cell 27, 137–152 (2016).

25. C. Makhoul et al., Intersectin-1 interacts with the golgin GCC88 to couple the actin network and Golgi architecture. Molecular biology of the cell 30, 370–386 (2019).

26. J. Bailey Blackburn, I. Pokrovskaya, P. Fisher, D. Ungar, V. V. Lupashin, COG Complex Complexities: Detailed Characterization of a Complete Set of HEK293T Cells Lacking Individual COG Subunits. Frontiers in cell and developmental biology 4, 23 (2016).

27. D. E. Gordon, L. M. Bond, D. A. Sahlender, A. A. Peden, A targeted siRNA screen to identify SNAREs required for constitutive secretion in mammalian cells. Traffic (Copenhagen, Denmark) 11, 1191–1204 (2010).

28. S. Liu, B. Storrie, How Rab proteins determine Golgi structure. International review of cell and molecular biology 315, 1–22 (2015).

29. F. Ferraro et al., A two-tier Golgi-based control of organelle size underpins the functional plasticity of endothelial cells. Developmental cell 29, 292–304 (2014).

30. J. J. McCormack, M. Lopes da Silva, F. Ferraro, F. Patella, D. F. Cutler, Weibel-Palade bodies at a glance. Journal of cell science 130, 3611–3617 (2017).

31. W. W. Lui-Roberts, L. M. Collinson, L. J. Hewlett, G. Michaux, D. F. Cutler, An AP-1/clathrin coat plays a novel and essential role in forming the Weibel-Palade bodies of endothelial cells. The Journal of cell biology 170, 627–636 (2005).

32. F. Ferraro et al., Weibel-Palade body size modulates the adhesive activity of its von Willebrand Factor cargo in cultured endothelial cells. Sci Rep 6, 32473 (2016).

33. L. Yuan et al., A role of stochastic phenotype switching in generating mosaic endothelial cell heterogeneity. Nature communications 7, 10160 (2016).

34. J. J. McCormack, K. J. Harrison-Lavoie, D. F. Cutler, Human endothelial cells size-select their secretory granules for exocytosis to modulate their functional output. J Thromb Haemost 18, 243–254 (2020).

35. M. Lopes da Silva, D. F. Cutler, von Willebrand factor multimerization and the polarity of secretory pathways in endothelial cells. Blood 128, 277–285 (2016).

36. T. D. Nightingale, K. Pattni, A. N. Hume, M. C. Seabra, D. F. Cutler, Rab27a and MyRIP regulate the amount and multimeric state of VWF released from endothelial cells. Blood 113, 5010–5018 (2009).

37. N. B. Cole, N. Sciaky, A. Marotta, J. Song, J. Lippincott-Schwartz, Golgi dispersal during microtubule disruption: regeneration of Golgi stacks at peripheral endoplasmic reticulum exit sites. Molecular biology of the cell 7, 631–650 (1996).

38. R. D. Starke et al., Cellular and molecular basis of von Willebrand disease: studies on blood outgrowth endothelial cells. Blood 121, 2773–2784 (2013).

39. F. Ferraro et al., Modulation of endothelial organelle size as an antithrombotic strategy. J Thromb Haemost 10.1111/jth.15084 (2020).

40. R. Ketteler et al., Image-based siRNA screen to identify kinases regulating Weibel-Palade body size control using electroporation. Scientific data 4, 170022 (2017).

41. M. Lopes-da-Silva et al., A GBF1-Dependent Mechanism for Environmentally Responsive Regulation of ER-Golgi Transport. Developmental cell 49, 786–801 e786 (2019).

42. N. L. Stevenson et al., G protein-coupled receptor kinase 2 moderates recruitment of THP-1 cells to the endothelium by limiting histamine-invoked Weibel-Palade body exocytosis. J Thromb Haemost 12, 261–272 (2014).

43. V. Kondylis, H. E. van Nispen tot Pannerden, B. Herpers, F. Friggi-Grelin, C. Rabouille, The golgi comprises a paired stack that is separated at G2 by modulation of the actin cytoskeleton through Abi and Scar/WAVE. Developmental cell 12, 901–915 (2007).

44. Y. Ravichandran, B. Goud, J. B. Manneville, The Golgi apparatus and cell polarity: Roles of the cytoskeleton, the Golgi matrix, and Golgi membranes. Curr Opin Cell Biol 62, 104–113 (2020).

45. S. Yadav, S. Puri, A. D. Linstedt, A primary role for Golgi positioning in directed secretion, cell polarity, and wound healing. Molecular biology of the cell 20, 1728–1736 (2009).

46. B. Bisel et al., ERK regulates Golgi and centrosome orientation towards the leading edge through GRASP65. The Journal of cell biology 182, 837–843 (2008).

47. H. C. Tie, A. Ludwig, S. Sandin, L. Lu, The spatial separation of processing and transport functions to the interior and periphery of the Golgi stack. Elife 7 (2018).

48. C. King, P. Sengupta, A. Y. Seo, J. Lippincott-Schwartz, ER membranes exhibit phase behavior at sites of organelle contact. Proceedings of the National Academy of Sciences of the United States of America 117, 7225–7235 (2020).

49. N. B. S. Cole, C.L.; Sciaky, N.; Terasaki, M.; Edidin, M.; Lippincott-Schwartz, J., Diffusional mobility of Golgi proteins in membranes of living cells. Science (New York, N.Y 273, 797–801 (1996).

50. H. L. Zenner, L. M. Collinson, G. Michaux, D. F. Cutler, High-pressure freezing provides insights into Weibel-Palade body biogenesis. Journal of cell science 120, 2117–2125 (2007).

## SI References

1. L. Hewlett et al., Temperature-dependence of Weibel-Palade body exocytosis and cell surface dispersal of von Willebrand factor and its propolypeptide. PLoS One 6, e27314 (2011).

2. F. Ferraro et al., A two-tier Golgi-based control of organelle size underpins the functional plasticity of endothelial cells. Developmental cell 29, 292–304 (2014).

3. H. L. Zenner, L. M. Collinson, G. Michaux, D. F. Cutler, High-pressure freezing provides insights into Weibel-Palade body biogenesis. Journal of cell science 120, 2117–2125 (2007).

